# Orthotopic Transplantation and Engraftment of Human Induced Pluripotent Stem Cell-Derived Alveolar Progenitor Cells into Murine Lungs

**DOI:** 10.1101/2020.06.14.149831

**Authors:** Aaron I. Weiner, Rafael Fernandez, Gan Zhao, Gargi Palashikar, Maria Fernanda de Mello Costa, Stephanie Adams, Christopher J. Lengner, F. Brad Johnson, Andrew E. Vaughan

## Abstract

Humanized mice possessing human cells, tissues, or organ systems provide an unparalleled platform for preclinical studies in oncology, immunology, and infectious diseases. While the lungs are a vital organ subject to a wide variety of pathologies, exemplified by the ongoing COVID-19 pandemic, discrete differences in murine and human lungs can obfuscate interpretation of murine models of lung disease. Here we provide proof-of-concept methodology for the potential humanization of murine lungs via orthotopic transplantation of human NKX2.1+ progenitor cells and alveolar type 2 cells derived from induced pluripotent stem cells. We show that these cells engraft readily into highly immunocompromised mice after pharmacological injury with bleomycin, which presumably generates “space” for human cells to access denuded basement membrane and engraft. Transplanted cells stably retain their pulmonary lineage restriction and persist as superficially differentiated alveolar type 2 and type 1 cells. Future work should focus on strategies to promote xenorepopulation of most / all of the murine lung with human cells while retaining appropriate regio-specific epithelial differentiation and normal physiological function.

## MAIN

While animal models for human disease research are essential for the ultimate development of new and effective therapies, it is widely accepted that their utility is limited. Even very promising advances in animal models often fail to be recapitulated in patient settings, and inflammatory responses in mice have been shown to be quite discrepant from humans^1^. In response, researchers have recognized that severely immunocompromised mice such as NOD *scid* gamma (NSG and NOG) strains lacking adaptive immunity and bearing compromised innate immune responses can serve as hosts for human organs and tissues^2^,^3^. These strains allow for stable maintenance of human cell types and even whole organs for years, thus providing a powerful platform to address important biological questions in human cells *in vivo*.

By far the most tractable humanized mouse models to date are mice with human immune systems, especially NSG mice bearing transplanted human CD34+ hematopoietic stem cells which demonstrate robust multi-lineage persistence of human immune cell types^3^,^4^. In terms of solid organs, liver repopulation with cadaveric human hepatocytes in FAH^-/-^ mice on a similar immunodeficient background (Rag-2^null^/IL-2Rγ^null^) is remarkably effective, resulting in up to 90% of the recipient liver being composed of human hepatocytes and rescuing otherwise fatal fumarylacetoacetate hydrolase deficiency^5^. Similar xenotransplantation of iPSC-derived human pancreatic β cells has also been recently demonstrated^6^, as has transplantation of human intestinal organoids^7^. In the lung, one group recently demonstrated successful engraftment of human stem cell-derived airway epithelial cells^8^, and another group achieved engraftment of primary (donor lung) derived type 2 pneumocytes (AT2s)^9^ but to the best of our knowledge successful engraftment of human iPSC-derived alveolar progenitor cells (AT2s), has not been demonstrated.

We have previously demonstrated successful syngeneic engraftment of murine type 2 cells via intranasal inhalation after various injurious stimuli (influenza, bleomycin, acid instillation)^9^. Assuming that human cells would be immediately rejected upon transplant into immunocompetent animals, we administered the chemotherapeutic agent bleomycin to NSG mice^3^ followed by orthotopic iPSC-derived lung/alveolar progenitor cell transplantation 10 days later. To generate donor cells, we utilized the well-established NGST BU3 line, which possesses fluorescent reporters for the lung epithelial transcription factor NKX2.1 and the AT2 marker surfactant protein C (SPC) (hereafter NKX2.1-GFP/SPC-tdTomato; NGST). These fluorescent markers allow for high-purity isolation of unspecified lung epithelial progenitors (GFP^+^ tdTomato^−^, hereafter NKX2.1^+^ epithelial progenitors) and AT2s (GFP^+^ tdTomato^+^; hereafter iAT2s) upon directed differentiation^10–12^. We transplanted either purified iAT2s or a mixture of NKX2.1^+^ lung progenitor cells, of which approx. 20% are SPC-tdTomato^+^ iAT2s at the time of transplant (Fig. 1A-B, Fig. 2). Recipient mice were euthanized 10-14 days post-transplant. In every transplant recipient examined (n=5) we could observe discrete clusters of cells present in the alveoli expressing several human antigens (NuMA, human-specific mitochondrial marker 113-1) as well as GFP, indicative of retention of the NKX2.1+ pulmonary lineage (Fig. 1C-G). We never observed any cells expressing human antigens but lacking GFP expression, indicating that even if a small fraction of NKX2.1^negative^ cells persist in culture, they do not successfully engraft.

**Figure 1.**
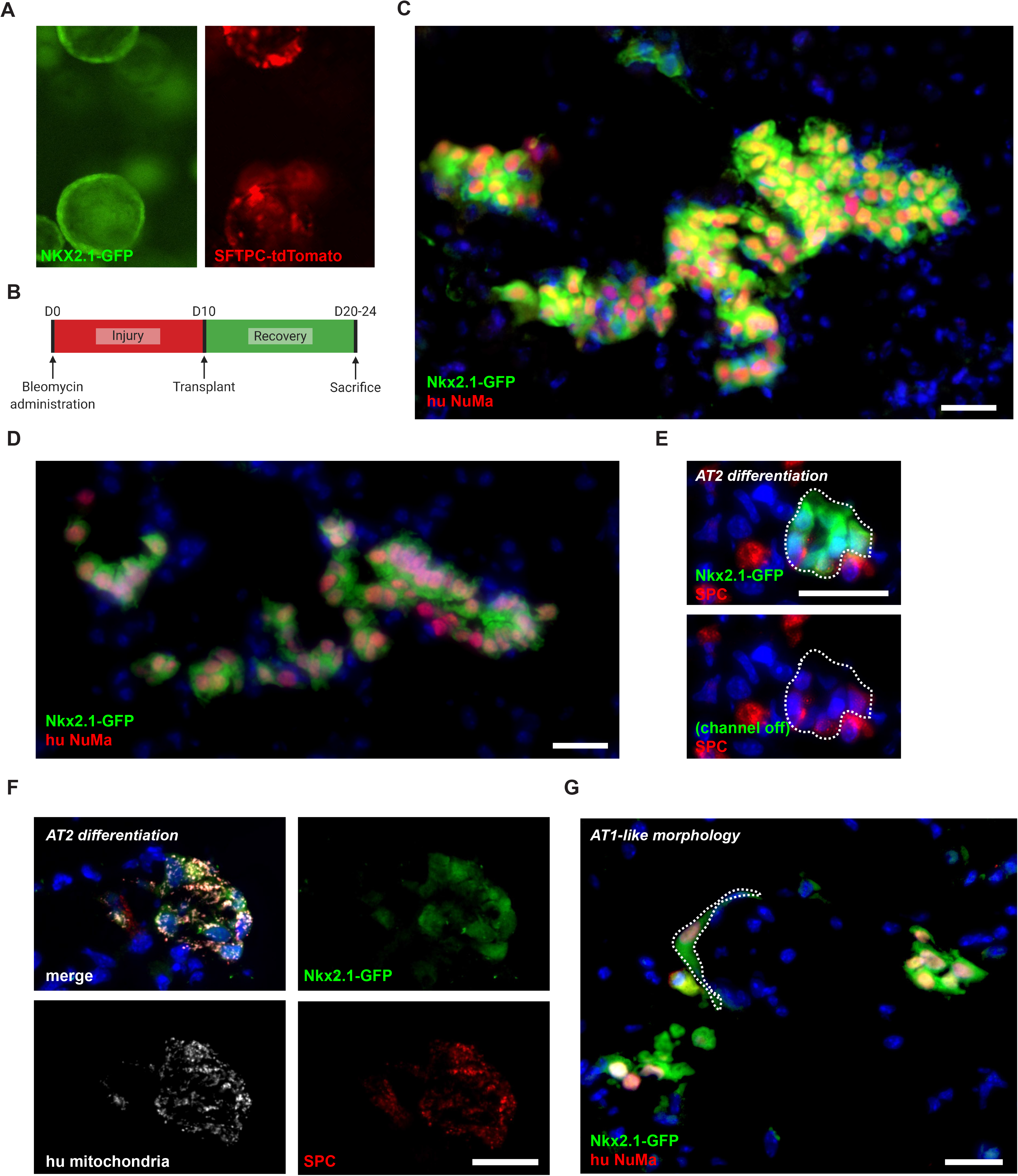
Human iPSC-derived lung progenitor cells engraft in injured, immunodeficient mouse lungs. A, Single channels from a representative image of NKX2.GFP+ lung progenitors cells derived from the BU3 NGST iPSC line cultured in Matrigel prior to dissociation for transplant, demonstrating a fraction of these cells exhibiting SFTPC-tdTomato reporter expression. B, Transplant timeline. C-G, Discrete engraftments of cells as in (A) transplanted into bleomycin-injured NSG mice. Some engraftments exhibited expression of the alveolar type 2 cell marker SPC (E, F) while others demonstrated an alveolar type 1 cell-like morphology (G). Scale bars = 25 μm.

**Figure 2.**
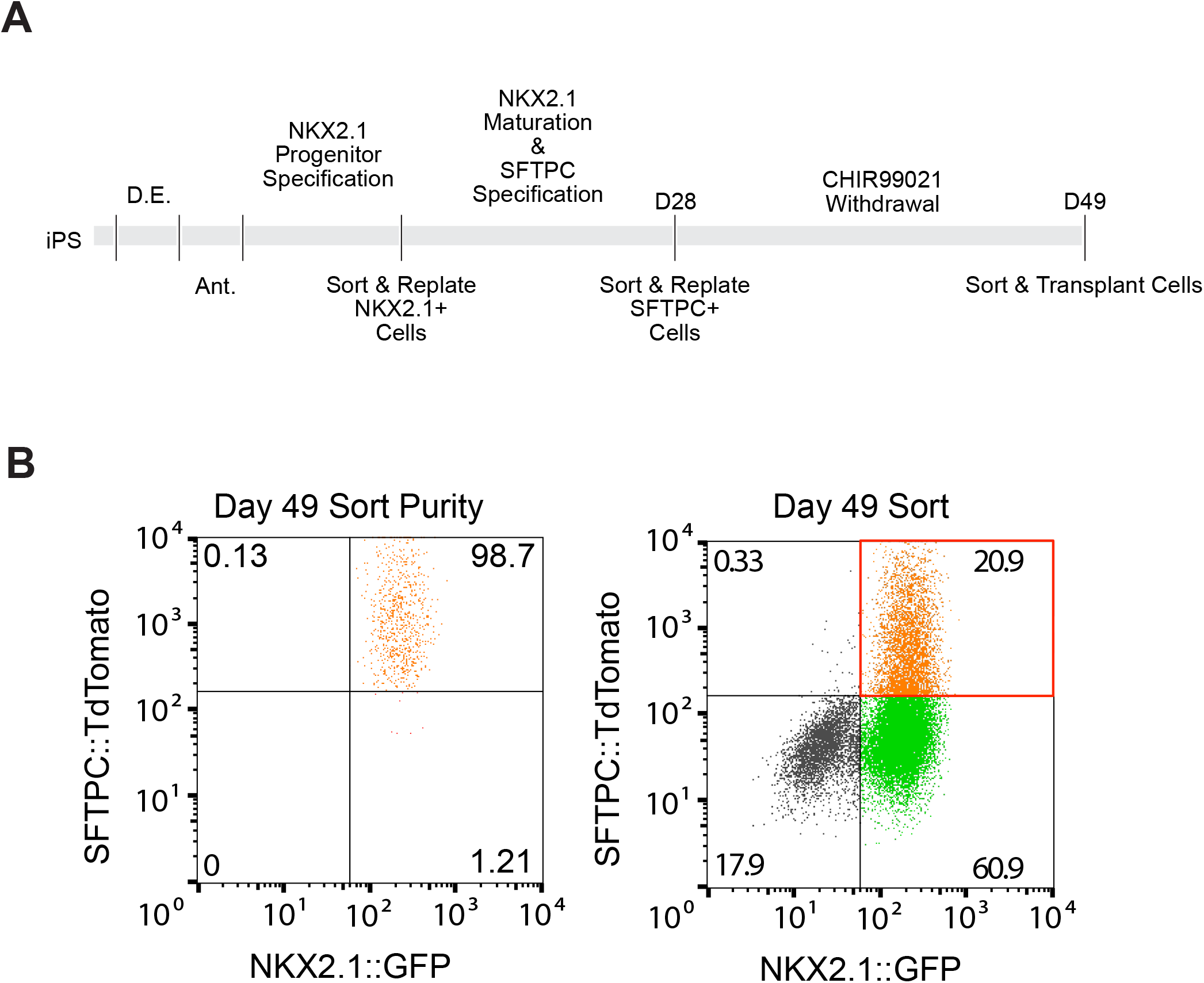
Differentiation scheme for human iPSCs (BU3 NGST) into NKX2.1+ lung progenitor cells and alveolar type 2 cells. A, Protocol for differentiation. B, Representative flow cytometry plots of cells isolated for transplantation after 49 days of differentiation.

We further explored the cell fates of transplanted cells. We observed many engrafted cells expressing SPC, evidencing either the retention of an AT2-like fate from the iAT2s already present in donor cultures or further *in vivo* differentiation of NKX2.1+ progenitors (Fig. 1E-F). We also observed some cells with morphological characteristics of alveolar type 1 cells (Fig. 1G), including a flattened, elongated morphology. We further stained for club cell and basal cell markers CC10 and Krt5, but did not observe expression of these markers in any of the transplanted cells. This is in contrast to a recent preprint demonstrating basal cell transdifferentiation of primary human AT2s derived from donor lungs^13^, though this discrepancy could be due to the fundamental differences in donor cells sources (primary tissue vs. iPSC-derived) or may require a larger sample size to observe.

While these results are preliminary and largely descriptive, this early proof-of-concept report opens major possibilities for future studies in pulmonary disease. For instance, SARS-CoV-2 cannot readily infect mice due to sequence dissimilarities in the ACE2 receptor^14^. Mice with both fully humanized lungs and immune systems would be expected to not only be readily infectable but could also more faithfully reproduce the pathophysiology of COVID-19 than in native mouse lungs. This platform also allows for *in vivo* analysis of iPSC-derived lung epithelial cells, especially those edited to correct or induce a disease state, the phenotypes of which can be difficult to model *in vitro*. This work is in very early stages, but it demonstrates exciting foundational data that generation of a humanized mouse lung may be possible in the near future.

## METHODS

### Animals

8-to 10-week-old mice were used for all experiments with males and females in roughly equal proportions. Experimenters were not blinded to mouse age or sexes. NOD.Cg-*Prkdc^scid^ Il2rg^tm1Wjl^/SzJ* (referred to as NSG) mice were utilized as recipients for all transplantation experiments. All animal experiments were carried out under the guidelines set by the University of Pennsylvania’s Institutional Animal Care and Use Committees and followed all NIH Office of Laboratory Animal Welfare regulations.

### Bleomycin injury

Mice were first anesthetized using 3.5% isoflurane in 100% O_2_ via an anesthesia vaporizer system. Mice were intranasally administered 4mg/kg body weight bleomycin sulfate (13877-10, Cayman Chemicals) in a total volume of 30 μL PBS. Only injured mice that lost ≥10% of their starting body weight by day 4 post-injury and survived to the time of transplant were considered to be adequately injured and used for all experiments involving bleomycin injury. Cells were transplanted at day 10 post-bleomycin administration.

### iPS Cell Line Maintenance

The BU3 NGST line was a generous gift from Dr. Darryl Kotton at Boston University. iPS cells used for differentiation were maintained on growth factor reduced Matrigel (Corning) coated plates in StemMACS™ iPS-Brew XF medium (Miltenyi Biotec). Cells were cultured in clusters and passaged every 4-5 days using StemMACS Dissociation reagent (Miltenyi Boitec) and passaged at a ratio of 1:6 to 1:12 depending on the cell line. All cells were routinely screened for mycoplasma contamination using a PCR based assay.

### Directed differentiation into NKX2.1+ lung progenitors and SFTPC+ iPS-derived AT2 cells

A modified version of the protocol described in Jacob et al. was used to generate SFTPC expressing iAEC2 cells. In brief, iPS cells were seeded at 500,000 cells per well on a 6-well plate with rock inhibitor for 24 hours and incubated at 5%O_2_ | 5% CO_2_ | 90% N2. Definitive endoderm was induced using the StemDiff Definitive Endoderm kit for 3 days. Next, the cells were split at a ratio of 1:3 onto fresh Matrigel plates and anteriorized using Dorsomorphin (2 μM) and SB431542 (10 μM) in cSFDM for three days. Cells were then differentiated into NKX2.1+ progenitors by incubating in CBRa media containing CHIR99021 (3 μM), BMP4 (10 ng/mL), and Retinoic Acid (100 nM) for 7 days changing media every 2 days at first and then increasing to every day media changes when the media became more acidic. On day 15 or 16, NKX2.1+ progenitors were sorted using a FACSJazz sorter using the endogenous NKX2.1::GFP reporter.

NKX2.1+ sorted cells were replated at a density of 400,000 cells/mL in 90% Matrigel supplemented with 10% of CK+DCI + TZV media (3 μM CHIR99021, 10 ng/mL KGF, 100 nM Dexamethasone, 100μM 8Br-cAMP and 100 μM IBMX and 2 μM TZV) (from now on referred to as 90/10 Matrigel). The Matrigel droplets were allowed to cure at 37°C for 20-30 minutes and then overlaid with an appropriate amount of CK+DCI + TZV Media. These organoid-containing matrigel droplets were incubated at 37°C at 20% O_2_, 5% CO_2_ and 75% N2 (room air) for 14 days changing with fresh media every other day. On Day 28, the iAEC2 containing organoids were sorted on a FACSJazz sorter for SPC+ cells using the endogenous SFTPC::TdTomato reporter. These sorted SPC+ cells were replated at a concentration of 65,000 cells / mL in 90/10 Matrigel drops and grown in K+DCI + TZV at 37°C at 20% O_2_, 5% CO_2_ and 75% N2 (Room air) for 3 weeks changing media every other day. The cells were then sorted again as before and transferred in CK + DCI + TZV media to be transplanted.

### Orthotopic Transplantation via Intranasal Inhalation

NSG mice were used as transplant recipients for all experiments. Transplant recipients received between 800,000 and 2 million cells. Recipient mice were anesthetized with 3.5% isoflurane in 100% O_2_ via an anesthesia vaporizer system and were intranasally administered cells by pipetting 30uL single-cell suspension in PBS + 1% P/S onto the nostrils of anesthetized mice in visually confirmed agonal breathing.

### Lung tissue preparation for immunostaining

Following sacrifice via isoflurane overdose, lungs were inflated at a constant pressure of 25 cm H2O with 3.2% paraformaldehyde (PFA) for 30 minutes followed by incubation in 3.2% PFA for another 30 minutes at room temperature. Fixed lungs were then washed in multiple PBS washes over the course of 1 hour at room temperature, followed by an overnight incubation in 30% sucrose shaking at 4 °C, and then a 2 hour incubation in 15% sucrose 50% OCT compound (Fisher HealthCare) at room temperature. Finally, fixed lungs were embedded in OCT by flash freezing with dry ice and ethanol.

### Immunostaining

7 μm sections were cut on a Leica CM3050 S Research Cryostat (Leica Biosystems). Tissue sections were further fixed for 5 minutes in 3.2% PFA, rinsed three times with PBS, and blocked in blocking solution (PBS + 1% bovine serum albumin (Affymetrix) + 5% normal donkey serum (Jackson Immuno Research) + 0.1% Triton X-100 (Millipore Sigma)+ 0.02% sodium azide (Millipore Sigma)) for > 30 minutes. Slides were incubated in primary antibodies (listed below) in blocking solution overnight at 4 °C. Slides were then washed three times with PBS + 0.1% Tween-20 (Millipore Sigma) and subsequently incubated with secondary antibodies (listed below) for > 2 h at room temperature. Slides were then washed once more with PBS + 0.1% Tween-20 prior to incubation in 1 μM DAPI (Life Technologies) for 5 minutes, rinsed with PBS, and mounted with either Prolong Gold (Life Sciences) or Fluoroshield (Millipore Sigma). The following primary antibodies were used: rabbit anti-NuMa (1:500, Thermo Fisher Scientific, PA5-22285), rabbit anti-SPC (1:2000, Millipore), mouse anti-human mitochondrial marker (1:500, Millipore, MAB1273 clone 1113), and sheep anti-eGFP (1:500, Invitrogen, 10396164). The following secondary antibodies were used: Alexa Fluor™ 488-conjugated donkey anti-sheep (1:1000, Thermo Fisher Scientific), Alexa Fluor™ 568-conjugated donkey anti-rabbit (1:1000, Thermo Fisher Scientific), Alexa Fluor™ 647-conjugated donkey anti-rabbit (1:1000, Thermo Fisher Scientific), and CruzFluor™ 647-conjugated mouse IgG kappa binding protein (1:1000, Santa Cruz Biotechnology).

